# *N*^6^-methyladenosine modification of HIV-1 RNA evades RIG-I-mediated sensing to suppresses type-I interferon induction in monocytic cells

**DOI:** 10.1101/2020.11.04.368712

**Authors:** Shuliang Chen, Sameer Kumar, Nagaraja Tirumuru, Jennifer L. Welch, Lulu Hu, Chuan He, Jack T. Stapleton, Li Wu

**Author notes:** School of Basic Medical Sciences, Wuhan University, Wuhan, Hubei, China (Shuliang Chen). Corresponding author: (LW). These authors contributed equally to this work (Shuliang Chen, Sameer Kumar).

## Abstract

*N*^6^-methyladenosine (m^6^A) is a prevalent RNA modification that plays a key role in regulating eukaryotic cellular mRNA functions. RNA m^6^A modification is regulated by two groups of cellular proteins, writers and erasers that add or remove m^6^A, respectively. HIV-1 RNA contains m^6^A modifications that modulate viral infection and gene expression in cells. However, it remains unclear whether m^6^A modifications of HIV-1 RNA modulate innate immune responses in cells or HIV-1-infected individuals. Here we show that m^6^A modification of HIV-1 RNA suppresses the expression of antiviral cytokine type-I interferon (IFN-I) in human monocytic cells. Transfection of differentiated monocytic cells with HIV-1 RNA fragments containing a single m^6^A-modification significantly reduced IFN-I mRNA expression relative to their unmodified RNA counterparts. We generated HIV-1 with altered RNA m^6^A levels by manipulating the expression of the m^6^A erasers or pharmacological inhibition of m^6^A addition in virus-producing cells. RNA transfection and viral infection of differentiated monocytic cells demonstrated that HIV-1 RNA with decreased m^6^A levels enhanced IFN-I expression, whereas HIV-1 RNA with increased m^6^A modifications had opposite effects. Our mechanistic studies revealed that m^6^A of HIV-1 RNA escaped the RIG-I-mediated RNA sensing and activation of the transcription factors IRF3 and IRF7 that drive IFN-I gene expression. Moreover, RNA of peripheral blood mononuclear cells from HIV-1 viremic patients showed increased m^6^A levels that correlated with increased *IFN-I* mRNA expression compared to levels from HIV-1-suppressed patients on antiretroviral therapy. Together, our results suggest that RNA m^6^A modifications regulate viral replication and antiviral innate immune responses in HIV-1-infected individuals.

**Author Summary:** HIV-1 is known as a weak inducer of antiviral cytokines including IFN-I, but it is unclear how HIV-1 evades innate immunity. Different types of RNA modifications including m^6^A within the HIV-1 genome modulate viral replication; however, the role of m^6^A modifications of HIV-1 RNA in regulating innate immune responses remains elusive. In this study, we found that HIV-1 RNA modified with m^6^A suppresses the expression of IFN-I in differentiated monocytic cells by avoiding innate immune detection of viral RNA mediated by RIG-I, an RNA sensor in host cells. We also observed significantly increased RNA m^6^A modifications of peripheral blood mononuclear cells from HIV-1 viremic patients compared to virally suppressed patients on combined antiretroviral therapy, suggesting a functional link between m^6^A modifications and antiretroviral treatment. Investigating the functions of m^6^A modifications of HIV-1 RNA in regulating innate immune sensing and IFN-I induction in monocytic cells can help understand the mechanisms of HIV-1 persistence.

## Introduction

Transcriptional modification of RNA in cells plays a crucial role in its stability, transportation, processing and thus regulation of gene expression. There are more than 160 RNA modifications identified in eukaryotes [1]. Methylation at the *N*^6^ position of adenosine (m^6^A) is a post-transcriptional RNA modification in internal and untranslated regions (UTRs) of eukaryotic mRNAs, microRNAs, small nuclear RNAs and long noncoding RNAs, which is important for RNA localization, stability and protein translation [1–5]. This methylation is controlled by two types of protein factors in cells, comprised of the writer complex [(methyltransferase-like 3 (METTL3) and METTL14] to incorporate methylation, and the erasers [fat mass and obesity associated protein (FTO) and α-ketoglutarate dependent dioxygenase AlkB homolog 5 (ALKBH5)] to remove m^6^A modification [6–9]. RNA m^6^A modification has been discovered in several RNA and DNA viruses over the past 40 years, although its effects on the viral lifecycle remain not fully understood [10–15]. Recent advancement of RNA sequencing based strategies expanded the identification and characterization of m^6^A to several clinically significant human pathogens [16], including HIV-1 [17–19]. Increasing evidence suggests that m^6^A modification plays a major role in regulation of viral replication and gene expression [16] and the immune system [20].

In the early stage of virus infections, sensing viral nucleic acids in infected cells is a critical step to induce innate immune responses that can lead to production of antiviral cytokines, including IFN-I (mainly IFN-α and IFN-β) [21]. Genomic RNA of HIV-1 and other viruses can be detected by cytosolic sensors, including retinoic acid-induced gene I (RIG-I) and melanoma differentiation-associated gene 5 (MDA5) [21]. Detection of viral RNA by these sensors triggers activation of several cellular kinases, which phosphorylate interferon regulatory factors 3 and 7 (IRF3 and IRF7) to induce IFN-I expression [22, 23]. HIV-1 is a weak inducer of host innate immune responses [24, 25], and it evades immune recognition by direct targeting of immune pathways, interacting with cellular proteins, or masking the viral genome from the cytosolic sensors [24, 26, 27]. HIV-1 RNA can be sensed by both RIG-I and MDA5, whereas it has evolved multiple strategies to escape innate immune surveillance [23, 28]. A recent study showed that 2’-*O*-methylation in HIV-1 RNA prevents MDA5-mediated sensing in myeloid cells, and thereby reduces IFN-I induction [29]. However, the role of m^6^A in regulating innate immune responses to HIV-1 RNA and the underlying mechanisms have not been defined.

Our previous *in vitro* studies showed that HIV-1 infection or HIV-1 envelope protein treatment of CD4^+^ T cells significantly up-regulates m^6^A levels of cellular RNA independently of viral replication [30]. However, it remains unclear whether m^6^A levels and IFN-I expression in HIV-1-infected individuals can be altered by effective antiretroviral therapy (ART), which leads to undetectable viral load in the vast majority of treated HIV-1 patients [31]. To address these fundamental questions and to better understand the role of m^6^A in HIV-1 infection *in vivo*, we measured the levels of m^6^A and IFN-I expression in peripheral blood mononuclear cells (PBMCs) of healthy donors, HIV-1 viremic patients before ART, and HIV-1 patients on ART.

Here we show that m^6^A modifications of HIV-1 RNA reduce viral RNA sensing and the induction of IFN-I in differentiated monocytic cells. We found that m^6^A-defective HIV-1 RNA induced IFN-I expression through RIG-I-mediated pathway, suggesting that m^6^A is an immune evasion strategy of HIV-1. In contrast to *in vitro* results, we also observed significantly increased levels of m^6^A RNA modifications and IFN-I expression in PBMCs from HIV-1 viremic patients compared to patients on ART. These results implicate that RNA m^6^A modifications can contribute to regulation of viral replication, innate immune responses, and ART in HIV-1-infected individuals.

## Results

### A single m^6^A modification of HIV-1 RNA oligos inhibits IFN-I induction in U937 cells

To examine the effect of m^6^A modification of HIV-1 RNA on IFN-I induction, we designed two different RNA oligos corresponding to two fragments of HIV-1 genome each with or without a single m^6^A modification [32] for transfection experiments. We have reported that these two m^6^A modifications in the 5’ untranslated regions (UTR) of HIV-1 genome are important for HIV-1 RNA binding to the m^6^A reader proteins (YTH domain family proteins 1-3) *in vitro* and viral replication in cells [32]. The m^6^A-modified RNA oligos 1 and 2 (both 42 mer) contained a single m^6^A-modified adenosine in the conserved GG**A**CU motif of the HIV-1 (NL4-3 strain) genome [32]. The m^6^A modification of the oligos was confirmed by immunoblotting with equal amounts of RNAs using m^6^A-specific antibodies (Fig. 1A and 1D). To mimic cellular responses to viral RNA in non-dividing macrophages, we differentiated monocytic U937 cells with phorbol 12-myristate 13-acetate (PMA) before transfection with the RNA oligos. Compared to unmethylated control (Ctrl) RNA, m^6^A-modified RNA oligo 1 induced 3-to 4-fold lower (*P* < 0.005) levels of *IFN-α* and *IFN-β* mRNA in transfected cells (Fig. 1B and 1C). Similar results were obtained with transfection of oligo 2, although the effects were less significant compared to oligo 1 (Fig. 1E and 1F). These results indicate that m^6^A modification of the 5’ UTR of HIV-1 RNA fragments inhibits IFN-I induction in differentiated U937 cells.

**Fig. 1.**
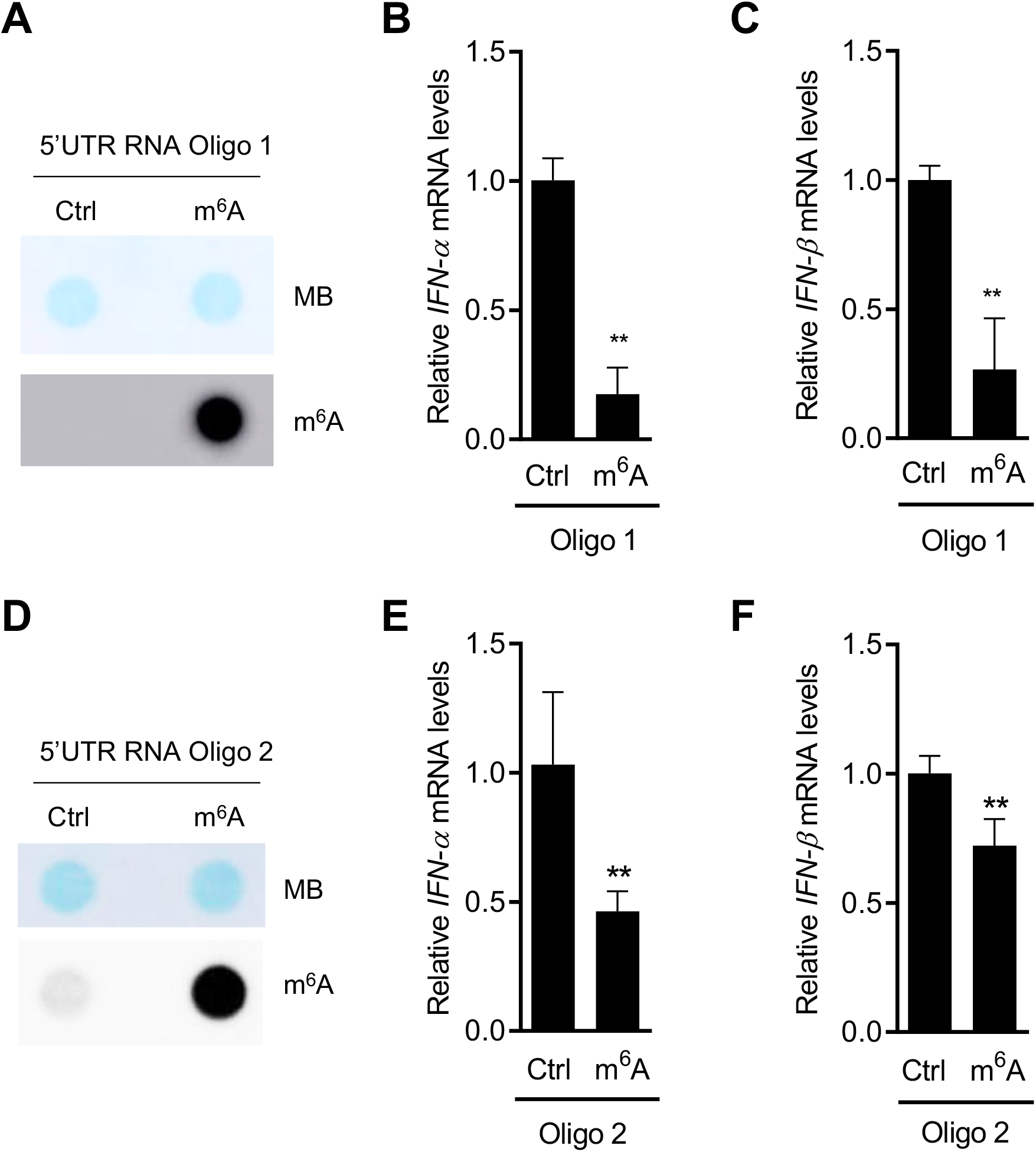
A single m^6^A modification of HIV-1 RNA oligos inhibits IFN-I induction in differentiated U937 cells. (**A**) HIV-1 5’ UTR (nt. 235–281) RNA oligo 1 (50 ng) with (m^6^A) or without (control, Ctrl) m^6^A modification were subjected to m^6^A dot-blot analysis. MB, methylene blue staining (an RNA loading control). (**B**) and (**C**) RNA oligo 1 (250 ng) were transfected into PMA-differentiated U937 cells. After 16 h, *IFN-α* and *IFN-β* mRNA levels were measured by RT-qPCR. Data shown are means ± S.D. of three independent experiments. Mann-Whitney t-test was used for statistical analysis. (**D**) HIV-1 5’ UTR (nt. 176–217) RNA oligo 2 (200 ng) with (m^6^A) or without (Ctrl) m^6^A modification were subjected to m^6^A dot-blot analysis. **(E)** and **(F)** RNA oligo 2 (250 ng) were transfected into PMA-differentiated U937 cells. After 16 h, *IFN-α* and *IFN-β* mRNA levels were measured by RT-qPCR. Data shown are means ± S.D. of three independent experiments. Un-paired t-test was used for statistical analysis. ** *P* < 0.005, compared with Ctrl samples.

### Inhibition of m^6^A modifications of HIV-1 RNA by FTO increases IFN-I induction

The m^6^A erasers (FTO and ALKBH5) orchestrate cellular mRNA functions by removing m^6^A modifications on mRNA [2]. To investigate whether m^6^A modifications of HIV-1 genomic RNA could suppress IFN-I induction in cells, purified RNA from HIV-1 virions was demethylated with recombinant FTO *in vitro*, resulting in a 10-fold decrease in m^6^A level relative to control HIV-1 RNAs (Fig. 2A). Transfection of m^6^A-reduced HIV-1 RNA into U937 cells induced 3-fold higher *IFN-α* and *IFN-β* expression (*P* < 0.0005) compared to control RNAs (Fig. 2B and 2C), suggesting that m^6^A modification of HIV-1 genomic RNA suppresses IFN-I induction in myeloid cells.

**Fig. 2.**
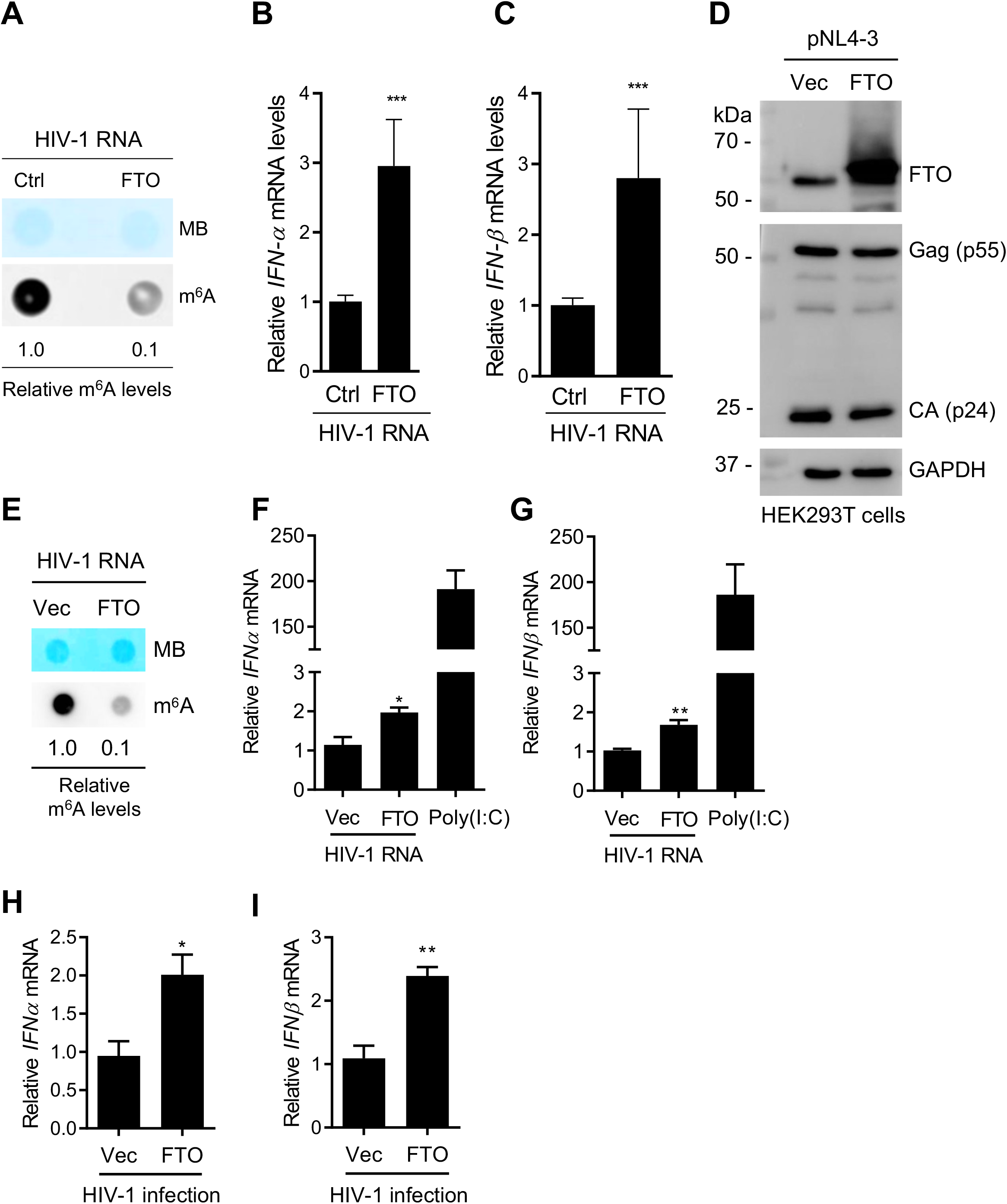
Inhibition of m^6^A modifications of HIV-1 RNA by FTO increases IFN-I induction. **(A)** m^6^A levels of HIV-1 genomic RNA were reduced by treatment with demethylase FTO and 50 ng of RNAs were used to confirm the m^6^A levels by the dot-blot assay. **(B)** and **(C)** 250 ng of the above RNAs were transfected in PMA-differentiated U937 cells. After 16 h, *IFN-α* and *IFN-β* mRNA levels were measured by RT-qPCR. The results are shown as means ± S.D. of three independent experiments. Mann-Whitney t-test was used for statistical analysis. *** *P* < 0.0005, compared with control samples. **(D)** HEK293T cells were transfected with vector control (Vec) or an FTO-expressing plasmid (FTO). After 24 h, HIV-1 proviral DNA clone (pNL4-3) was transfected for 48 h. Then, cell lysates were collected, and Western blotting was performed using indicated antibodies. **(E)** the m^6^A levels in HIV-1 genomic RNA were determined by the dot-blot assay using 100 ng purified viral RNA derived from Vec or FTO-expressing HEK293T cells. (**F**) and **(G)** PMA-differentiated U937 cells were transfected with 500 ng of HIV-1 RNA or 250 ng of poly(I:C) for 16 h and analyzed for *IFN-α* and *IFN-β* mRNA levels by RT-qPCR. The results are shown as means ± S.D. of three independent assays. Un-paired t-test was used for statistical analysis. * *P* < 0.05, ** *P* <0.005, compared between FTO and Vec samples. **(H)** and **(I)** PMA-differentiated U937 cells were infected by HIV-1 (250 pg of p24) from Vec or FTO-expressing HEK293T cells for 16 h, and *IFN-α* and *IFN-β* mRNA levels were quantified by RT-qPCR. The results are shown as means ± S.D. of three independent assays. Un-paired t-test was used for statistical analysis. * *P* < 0.05, ** *P* <0.005, Vec samples were normalized with non-infection samples.

To determine the effect of m^6^A of HIV-1 RNA on IFN-I induction during viral infection, HIV-1 containing lower levels of m^6^A in viral RNA was generated by overexpression of the eraser FTO in HIV-1-producing HEK293T cells. Compared to the vector control, FTO overexpression in HEK293T cells had no significant effect on the expression of HIV-1 Gag and capsid (CA, or p24) proteins (Fig. 2D). HIV-1 derived from FTO-overexpressed HEK293T cells (m^6^A-lower HIV-1) showed 10-fold lower m^6^A levels of viral RNA compared to viruses derived from control cells (Fig. 2E). When PMA-differentiated U937 cells were transfected with RNA of m^6^A-lower HIV-1, a 2-fold increase (*P* < 0.05) of *IFN-α* and *IFN-β* expression was observed compared to control HIV-1 RNA (Fig. 2F and 2G). As a positive control, poly(I:C) induced approximately 190-fold increases of *IFN-α* and *IFN-β* expression in transfected U937 cells (Fig. 2F and 2G). Moreover, differentiated U937 cells infected with m^6^A-lower HIV-1 expressed 2-fold higher (*P* < 0.05) *IFN-α* and *IFN-β* relative to control HIV-1 (Fig. 2H and 2I). Thus, HIV-1 containing reduced RNA m^6^A modifications induces higher *IFN-I* expression in differentiated U937 cells.

### Inhibition of m^6^A modifications of HIV-1 RNA by ALKBH5 increases IFN-I induction

To confirm the results of FTO treatment and overexpression, we also examined the effect of another m^6^A eraser ALKBH5 on HIV-1 RNA-mediated IFN-I induction in PMA-differentiated U937 cells. ALKBH5 overexpression in HIV-1-producing HEK293T cells had no significant effect on the expression of HIV-1 Gag and CA (Fig. 3A). The m^6^A modification of HIV-1 RNA generated from ALKBH5-overexpressed HEK293T cells showed a 2-fold decrease compared to HIV-1 RNA from control cells (Fig. 3B). *IFN-α* and *IFN-β* levels in U937 cells transfected with HIV-1 RNA from ALKBH5-overexpressed HEK293T cells were 1.8-fold higher (*P* < 0.05) compared to that from control cells (Fig. 3C and 3D). Furthermore, infection of U937 cells with HIV-1 from ALKBH5-overexpressed HEK293T cells induced 2-fold higher *IFN-α* and *IFN-β* expression (*P* < 0.0005) compared to HIV-1 from control HEK293T cells (Fig. 3E and 3F). Thus, inhibition of m^6^A modifications of HIV-1 RNA by eraser overexpression in virus-producing cells increases *IFN-I* induction in differentiated U937 cells.

**Fig. 3.**
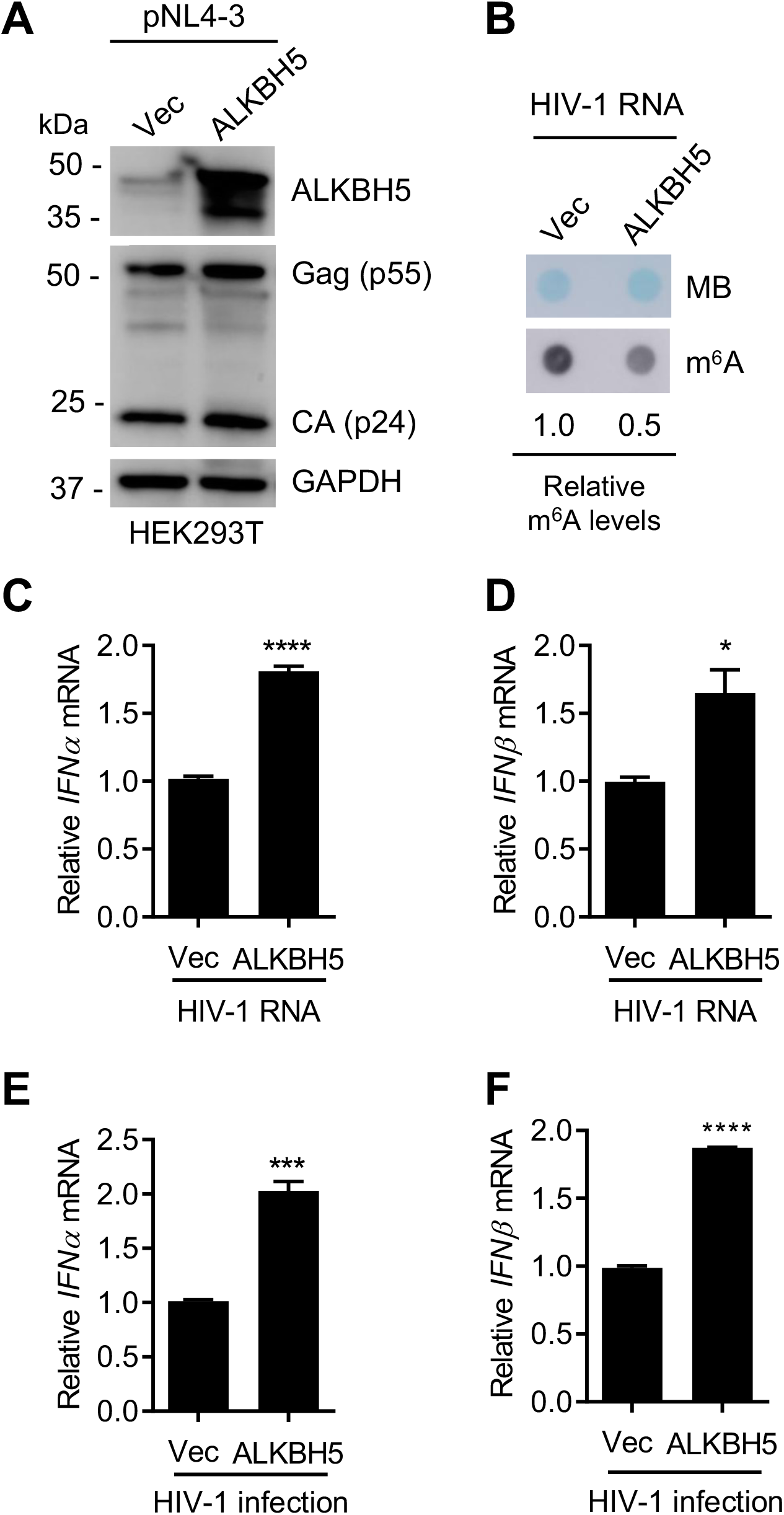
Inhibition of m^6^A modifications of HIV-1 RNA by ALKBH5 increases IFN-I induction. **(A)** HEK293T cells were transfected with a vector control (Vec) or an ALKBH5-expressing plasmid (ALKBH5). After 24 h, pNL4-3 was transfected into these cells for 48 h. Western blotting of cell lysates was performed using specific antibodies. **(B)** HIV-1 genomic RNA m^6^A levels were determined by the dot-blot assay using 100 ng viral RNA from Vec or ALKBH5-expressing HEK293T cells. **(C)** and **(D)** PMA-differentiated U937 cells were transfected with 500 ng of the indicated HIV-1 RNAs. At 16 h post-transfection, cells were collected for the analysis of *IFN-α* and *IFN-β* mRNA levels by RT-qPCR. The results are shown as means ± S.D. of three repeated assays. * *P* < 0.05, **** *P* <0.0001. **(E)** and **(F)** PMA-differentiated U937 cells were infected with HIV-1 (250 pg of p24) from Vec or ALKBH5-expressing HEK293T cells for 16 h, and *IFN-α* and *IFN-β* mRNA levels were quantified by RT-qPCR. The results are shown as means ± S.D. of three repeated experiments. Vec samples were normalized with non-infection samples. Un-paired t-test was used for statistical analysis. *** *P* < 0.0005, **** *P* <0.0001. Vec samples were normalized with non-infection samples. Un-paired t-test was used for statistical analysis.

### Knockout (KO) of erasers increases m^6^A levels in HIV-1 RNA and reduces IFN-I induction

To validate the results from eraser overexpression, we constructed FTO-KO and ALKBH5-KO HEK293T cell lines by the CRISPR-Cas9 method. Next, these cell lines were transfected to generate HIV-1 with increased m^6^A of viral RNA. Western blotting results showed that FTO and ALKBH5 were completely silenced and HIV-1 Gag protein expression was not significantly affected by FTO and ALKBH5 knockout (Fig. 4A). HIV-1 RNA from FTO-KO and ALKBH5-KO cells showed 7- and 25-fold higher m^6^A levels, respectively, relative to that from control (Con-KO) cells (Fig. 4B). Transfection of PMA-differentiated U937 cells with HIV-1 RNA derived from FTO-KO or ALKBH5-KO cells showed a 3-4-fold decrease (*P* < 0.05) in *IFN-I* expression compared to that from Con-KO cells (Fig. 4C and 4D). Moreover, infection of PMA-differentiated U937 cells with HIV-1 from FTO-KO or ALKBH5-KO cells induced approximately 2-fold less *IFN-I* expression (*P* <0.005) compared to Con-KO cells (Fig. 4E and 4F). Thus, increasing m^6^A levels in HIV-1 RNA by eraser KO in virus-producing cells reduces *IFN-I* induction in differentiated monocytic cells.

**Fig. 4.**
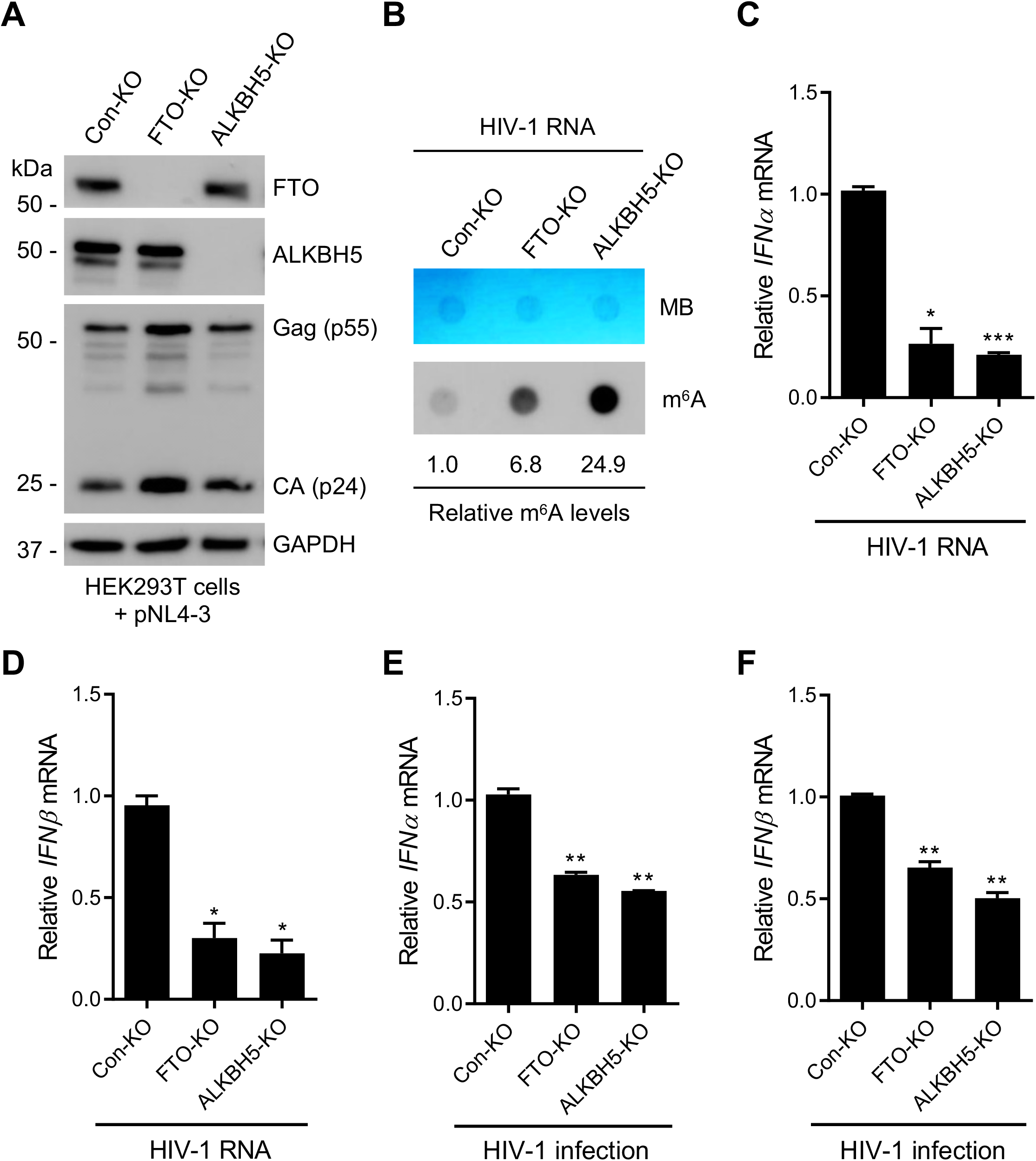
Knockout of erasers increases m^6^A levels in HIV-1 RNA and reduces IFN-I induction. **(A)** A single clone-derived control, FTO or ALKBH5 knockout (KO) HEK293T cells were transfected with pNL4-3 HIV proviral DNA. After 48 h, cells were collected for Western blotting analysis. **(B)** HIV-1 from the KO cells were collected and viral genomic RNA m^6^A level was determined by the dot-blot assay using 200 ng viral RNA. **(C)** and **(D)** HIV-1 RNA (250 ng) from KO cells were transfected into PMA-differentiated U937 cells. After 16 h, cells were collected for the analysis of *IFN-α* and *IFN-β* mRNA levels by RT-qPCR. The results are shown as means ± S.D. of three repeats with similar result. * *P* < 0.05, *** *P* < 0.0005. Un-paired t-test was used for statistical analysis. **(E)** and **(F)** HIV-1 (250 pg of p24) from KO cells were used to infect PMA-differentiated U937 cells for 16 h, and cells were collected for the analysis of *IFN-α* and *IFN-β* mRNA levels by RT-qPCR. The results are shown as means ± S.D. of three repeats with similar result. ** *P* <0.005. Un-paired t-test was used for statistical analysis.

### m^6^A-defective HIV-1 RNA induces IFN-I expression through IRF3 and IRF7 phosphorylation

We next investigated pharmacological inhibition of m^6^A modification using 3-deazaadenosine (DAA), an inhibitor of S-Adenosylhomocysteine (SAH) hydrolase that can catalyze the reversible hydrolysis of SAH to adenosine and homocysteine [33]. DAA causes SAH accumulation thereby elevating the ratio of SAH to S-adenosylmethionine (SAM), a substrate of m^6^A modification, and subsequent inhibition of SAM-dependent methyltransferases [33]. DAA-treatment of HEK293T cells did not affect HIV-1 production and release, but reduced m^6^A level in HIV-1 RNA 7-fold compared to control cells (Fig. 5A and 5B). Transfection of PMA-differentiated U937 cells with purified RNA from HIV-1 produced from DAA-treated HEK293T cells (DAA-HIV-1) induced 15-fold and 2.3-fold higher *IFN-α* and *IFN-β* expression (*P* < 0.0005), respectively (Fig. 5C and 5D). Moreover, infection of PMA-differentiated U937 cells with DAA-HIV-1 induced a 2-3-fold increase in *IFN-I* expression (*P* < 0.0005) compared to viruses from control HEK293T cells (Fig. 5E and 5F). These data further validate that m^6^A of HIV-1 RNA suppresses IFN-I induction in differentiated monocytic cells.

**Fig. 5.**
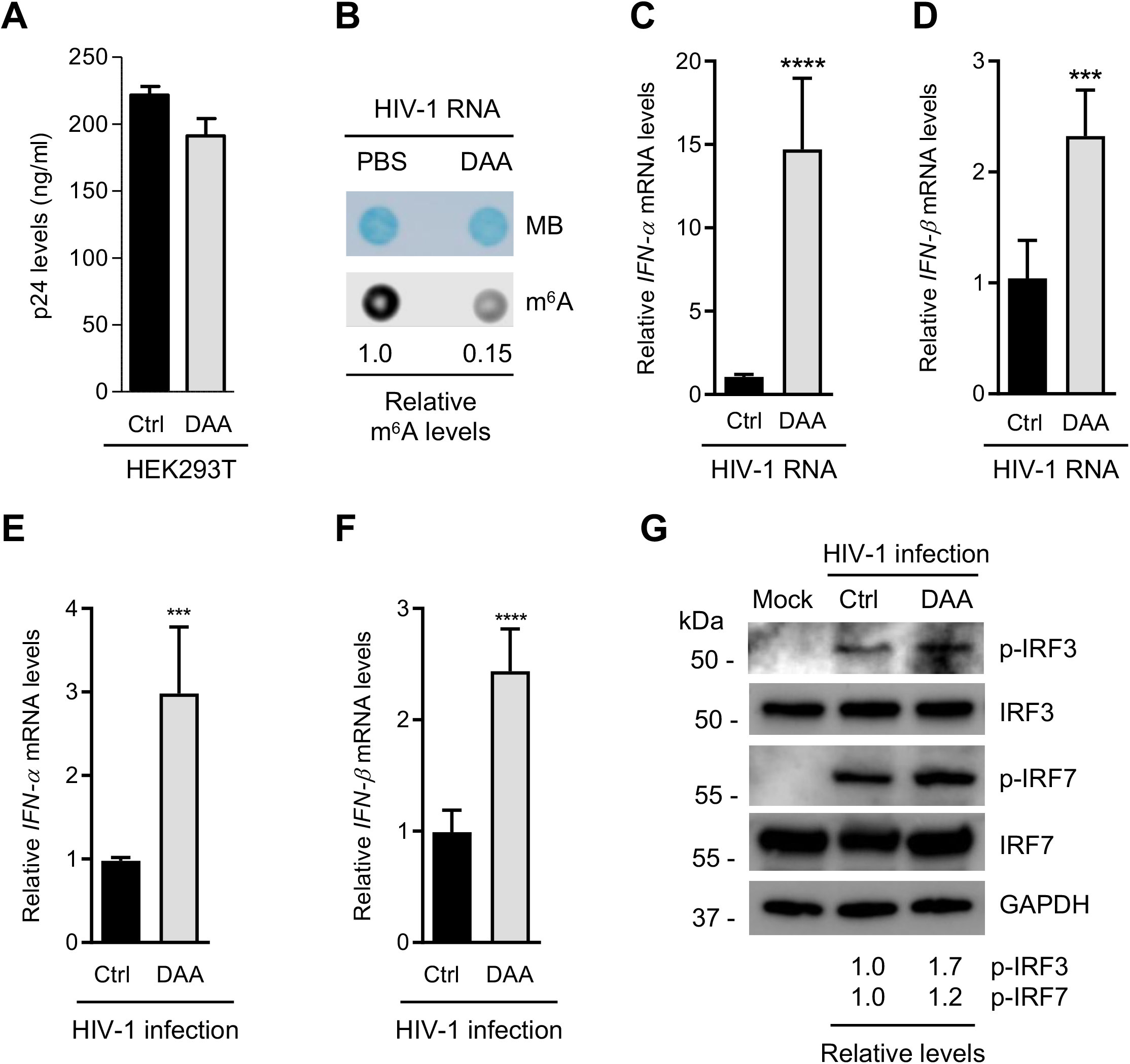
DAA-treatment reduces m^6^A modifications of HIV-1 RNA and increases IFN-I induction through IRF3 and IRF7 phosphorylation. HEK293T cells were treated with solvent (Ctrl, PBS) or DAA (50 μM) for 3 h and then transfected with the HIV-1 proviral DNA pNL4-3. HIV-1 in the supernatants was collected after 48 h. **(A)** HIV-1 p24 levels in the supernatants were measured by ELISA. **(B)** RNA (100 ng) from these viruses used for the m^6^A dot-blot assay. **(C)** and **(D)** HIV-1 RNA (250 ng) from Ctrl and DAA-treated samples were transfected into PMA-differentiated U937 cells. After 16 h, cells were collected for the analysis of *IFN-α* and *IFN-β* mRNA levels by RT-qPCR. The results are shown as means ± S.D. of three independent experiments. *** *P* < 0.0005, **** *P* < 0.0001. Un-paired t-test was used for statistical analysis. **(E)** and **(F)** HIV-1 (250 pg of p24) from HEK293T cells was used to infect PMA-differentiated U937 cells for 16 h. After 16 h, U937 cells were collected for the analysis of *IFN-I* mRNA levels by RT-qPCR. The results are shown as means ± S.D. of three independent experiments. *** *P* < 0.0005, **** *P* < 0.0001, Ctrl samples were normalized with non-infection samples. Un-paired t-test was used for statistical analysis. **(G)** PMA-differentiated U937 cells were infected with HIV-1 (250 pg of p24) derived from DAA-treated or control HEK293T cells for 4 h, and U937 cell lysates (50 μg proteins/sample) were used for the analysis of the indicated proteins by Western blotting. GAPDH is used as a loading control. The p-IRF3 and p-IRF7 indicate phosphorylated IRF3 and IRF7, respectively.

Because IFN-I expression is predominately driven by IRF3 and IRF7 after their activation by phosphorylation upon virus infections [21, 34], we tested whether DAA-HIV affected phosphorylation of IRF3 and IRF7. Compared to mock-infected U937 cells, control HIV-1 and DAA-HIV-1 induced strong phosphorylation of IRF3 and IRF7 in differentiated U937 cells at 4 h post-infection (Fig. 5G). Notably, phosphorylation of IRF3 and IRF7 was 1.7-fold and 1.2-fold higher in U937 cells infected with DAA-HIV-1 relative to control HIV-1, respectively (Fig. 5G). These results suggest that inhibition of HIV-1 RNA m^6^A modification triggers innate immune responses by inducing IRF3/7-mediated IFN-I expression in myeloid cells.

### RIG-I, but not MDA5, contributes to m^6^A modification of HIV-1 RNA induced IFN-I expression

To characterize the cellular sensing mechanisms of m^6^A-defective HIV-1 RNA, RIG-I and MDA5 in U937 cells were silenced by KO and shRNA, respectively. RIG-I-KO U937 cells were constructed and undetectable RIG-I expression was confirmed (Fig. 6A). To test whether these cells responded to RNA stimulation, poly(I:C) was transfected into cells and *IFN-I* expression was measured. Compared to untransfected cells (mock), poly(I:C) transfection induced high levels of *IFN-I* expression in RIG-I-KO and control U937 cells (Fig. 6B). As expected, the induction of *IFN-I* by poly(I:C) was significantly reduced by 2-fold in RIG-I-KO U937 cells (*P* < 0.005) compared to control cells (Fig. 6B), confirming that RIG-I acted as an RNA sensor to induce *IFN-I* expression in these cells. In control U937 cells, transfection of single m^6^A-modified HIV-1 RNA oligos induced lower *IFN-I* expression (*P* < 0.0001) compared to unmethylated RNA oligo counterparts (Fig. 6C and 6D). However, in RIG-I-silenced U937 cells, transfection of m^6^A-modified HIV-1 RNA oligos had no effect on *IFN-I* expression relative to unmethylated control oligos (Fig. 6C and 6D), suggesting a pivotal role of RIG-I in sensing m^6^A-defective HIV-1 RNA.

**Fig. 6.**
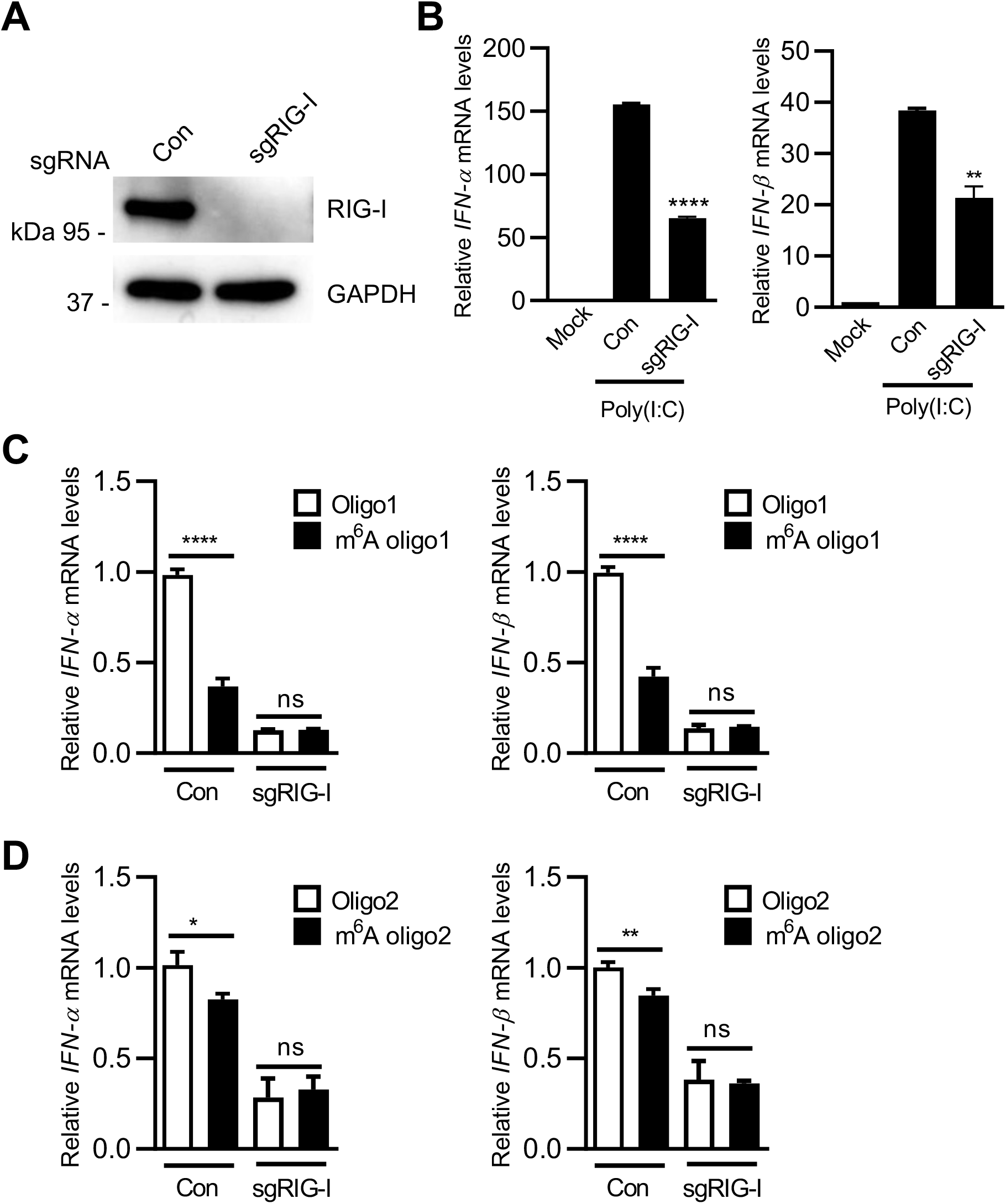
RIG-I senses m^6^A modification of HIV-1 RNA to induce IFN-I expression. **(A)** RIG-I expression levels in control (Con) and RIG-I knockout (sgRIG-I) U937 cells were measured by Western blotting. **(B)** Con and RIG-I KO U937 cells were transfected with 250 ng of poly(I:C). At 16 h post-transfection, cells were collected for the analysis of *IFN-α* and *IFN-β* mRNA levels by RT-qPCR. The results are shown as means ± S.D. of three repeats with similar result. ** *P* < 0.005, **** *P* < 0.0001. **(C)** and **(D)** PMA-differentiated Con and RIG-I KO U937 cells were transfected with 250 ng of RNA oligo 1 **(C)** or oligo 2 **(D)**. After 16 h, cells were collected for the analysis of *IFN-α* and *IFN-β* mRNA levels by RT-qPCR. The results are shown as means ± S.D. of three repeated experiments. * *P* < 0.05, ** *P* < 0.005, **** *P* < 0.0001. Un-paired t-test was used for statistical analysis. ns, not significant.

Furthermore, we examined the potential role of MDA5 in sensing m^6^A-defective HIV-1 RNA in monocytic cells. MDA5 expression was substantially reduced in differentiated U937 cells with MDA5 knockdown (shMDA5) compared to vector control (shCon) cells (Fig. 7A). As a positive control, poly(I:C) transfection induced high levels of *IFN-I* expression in both shCon and shMDA5 U937 cells. As expected, poly(I:C) transfection into shMDA5 U937 cells significantly decreased *IFN-I* levels relative to shCon cells (Fig. 7B). These cells were then examined for their ability to induce *IFN-I* expression by HIV-1 5’ UTR RNA oligos with or without single m^6^A modification [32]. Compared to unmethylated HIV-1 RNA oligos, transfection of m^6^A-modified HIV-1 RNA oligos reduced *IFN-I* expression in both shCon and shMDA5 U937 cells (Fig. 7C and 7D), suggesting that MDA5 is not a specific cellular sensor to detect m^6^A-defective HIV-1 RNA in differentiated monocytic cells.

**Fig. 7.**
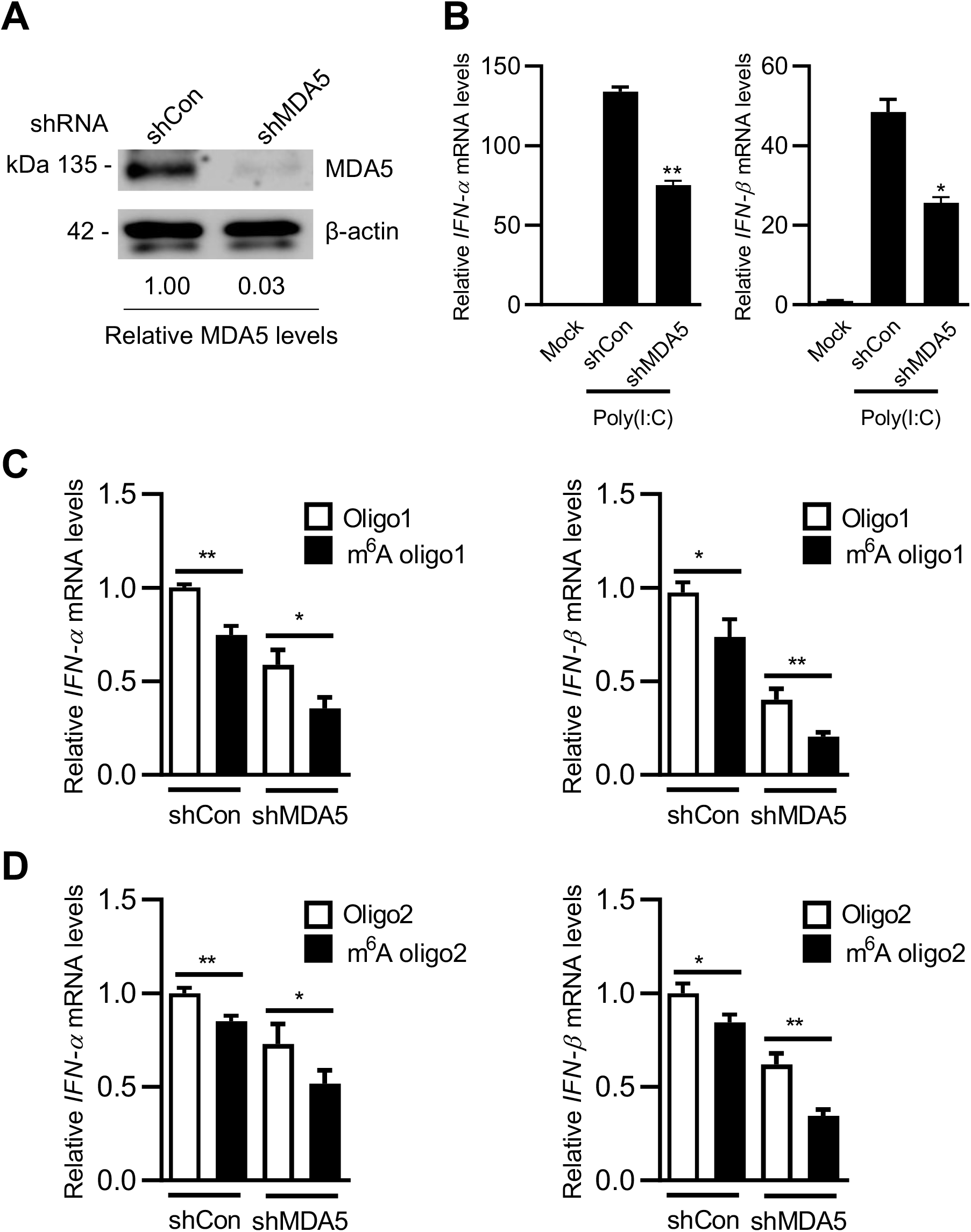
MDA5 has no specific role in m^6^A modification of HIV-1 RNA to induce IFN-I expression. **(A)** MDA5 expression levels were measured by Western blotting using control (shCon) and stable MDA5 knockdown (shMDA5) U937 cells. **(B)** shCon and shMDA5 U937 cells were transfected with poly(I:C). At 16 h post-transfection, cells were collected for the analysis of *IFN-α* and *IFN-β* mRNA levels by RT-qPCR. The results are shown as means ± S.D. of three repeats with similar result. * *P* < 0.05, ** *P* < 0.005. **(C)** and **(D)** PMA-differentiated shCon and shMDA5 U937 cells were transfected with 250 ng of RNA oligo 1 **(C)** or oligo 2 **(D)**. At 16 h post-transfection, cells were collected for the analysis of *IFN-α* and *IFN-β* mRNA levels by RT-qPCR. The results are shown as means ± S.D. of three repeated experiments.

### HIV-1 infected patients have higher level of m^6^A modification in the RNA of PBMCs

To explore the significance of m^6^A modifications in HIV-1-infected individuals and investigate the effect of ART on m^6^A levels, we measured the levels of m^6^A and *IFN-I* mRNA in immune cells from HIV-1 viremic patients in comparison with healthy control donors and HIV-1 patients on ART. We obtained PBMCs from healthy control donors (n=9), HIV-1 viremic patients (n=6) with different viral load pre-therapy, and HIV-1-infected individuals on ART (n=16) whose viral load was undetectable for a minimum of 6 months (<20 copies/mL) (supplemental Table S1). The average m^6^A level in total RNA of PBMCs from HIV-1 viremic patients was significantly higher (*P* < 0.005) compared to that from patients on ART (Fig. 8A and Supplemental Fig. S1A), suggesting an inverse correlation between viral load and m^6^A level of patient PBMCs. A visible, but not statistically significant increase (*P* = 0.27) in cellular RNA m^6^A level was observed in viremic patients compared to healthy individuals (Fig. 8A and Fig. S1A). This observation is also consistent with our previous results showing that HIV-1 infection or treatment of cells with HIV-1 envelope proteins (Env) upregulated m^6^A levels in primary CD4^+^ T-cells *in vitro* [30]. It is possible that Env shedding from HIV-1 viremic patients could upregulate m^6^A levels in PBMCs.

**Fig. 8.**
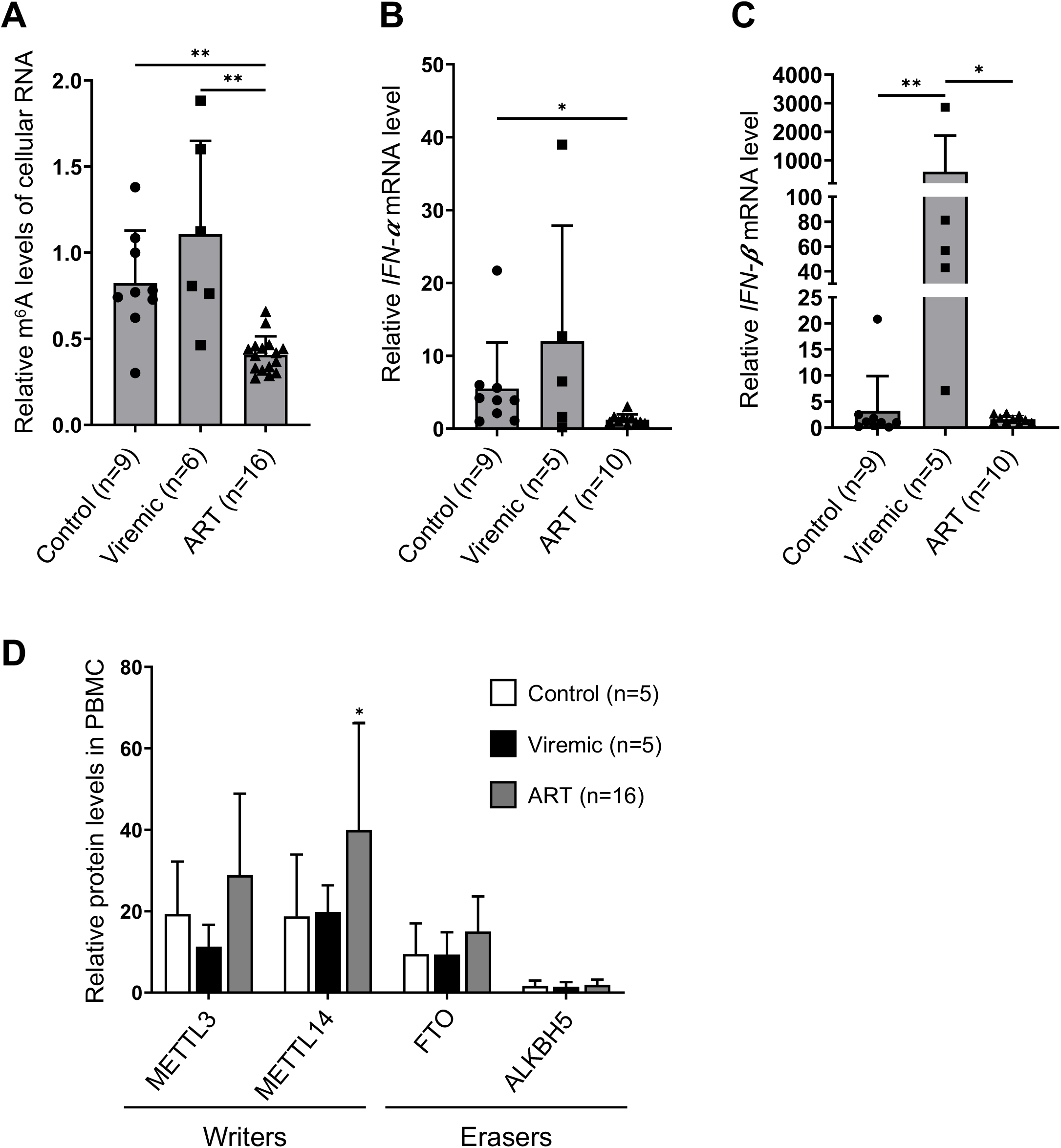
Increased m^6^A levels in total RNA of PBMCs from HIV-1 viremic patients. **(A)** Total cellular RNA was isolated from the PBMCs of uninfected healthy individuals (control), HIV-1 infected individuals without ART (viremic), and HIV-1 infected individuals on ART (ART) were subjected to m^6^A dot-blot analysis (200 ng RNA). The dots were quantified and normalized to the respective methylene blue control. The m^6^A level of first control sample (C1) was set as 1. Each symbol within the column represents each individual’s data point. **(B)** and **(C)** The isolated RNA from the PBMCs of the above described three groups of patients were subjected to RT-qPCR for measuring *IFN-α* and *IFN-β* mRNA expression. The values were normalized to their respective internal control (*GAPDH*). Each symbol represents the data from each individual. **(D)** Cell lysates of PBMCs of each group were subjected to Western blotting for m^6^A writers (METTL3 and METTL14) and erasers (ALKBH5 and FTO). The bands were quantified using Image J software and normalized to GAPDH control before plotting into this graphical representation. n, the number of healthy donors or patients. Five samples (C6-C9 and V6) were not included in the Western blot analysis due to the lack of sufficient cell lysates. * *P* < 0.05, ** *P* < 0.005. The one-way analysis of variance (ANOVA) nonparametric was used for statistical analysis.

Next, the levels of *IFN-I* mRNA in PMBCs were measured to analyze potential correlations with the RNA m^6^A levels. Compared to PBMCs from healthy donors, there was a trend of increase in *IFN-α* expression, and a significant increase in *IFN-β* expression in PBMCs from the viremic patients (Fig. 8B and 8C). Compared to PBMCs from the viremic patients, a significant decrease in both *IFN-α* and *IFN-β* expression (*P* < 0.005) was observed in patients on ART (Fig. 7B and 7C), suggesting that HIV-1 suppression by ART reduces innate immune responses to viral infection. Consistent with our results, a previous study [35] also reported similar results of decreased IFN-α expression in HIV-1 patients on ART compared to patients without ART.

Enhanced m^6^A levels in viremic patients could not be attributed to inherent differences in the levels of m^6^A writers and erasers, because there was no significant change in the expression of METTL3, FTO and ALKBH5 in PBMCs from the three groups (Fig. 8D and Fig. S1B). However, a slight increased level of METTL14 expression was observed in PBMCs from patients with ART compared to viremic patients (Fig. 8D and Fig. S1B). Together, these results suggest that HIV-1 infection upregulates the m^6^A level of cellular RNA in PBMCs from viremic patients without altering the expression of the writers and erasers.

## Discussion

HIV-1 genomic RNA contains 10-14 sites of m^6^A modifications in the 5’-, 3’-UTR and several coding regions [17–19]. Recent studies indicate that m^6^A modification has important effects on HIV-1 replication, gene expression, and host responses to viral infection [30, 32, 36].

It has been shown that cellular enzymes involved in RNA m^6^A modifications negatively regulate the innate immune response to infection of human cytomegalovirus, influenza A virus, adenovirus, or vesicular stomatitis virus by targeting the IFN-I pathway [37, 38]. A recent study showed that m^6^A modifications of human metapneumovirus RNA mimic the host RNA to avoid RIG-I-mediated innate immune sensing, and thereby reduce the production of IFN-I and enhance viral replication [39]. However, it remains unknown whether m^6^A modifications of HIV-1 RNA have any impact on innate immune responses.

In this study, we show that m^6^A modifications of HIV-1 RNA act as a negative regulator of IFN-I induction by avoiding RIG-I-mediated RNA sensing in PMA-differentiated U937 cells. We observed that two different HIV-1 RNA oligos of the HIV-1 5’-UTR containing a single m^6^A-modification significantly reduced IFN-I induction relative to their unmodified RNA counterparts. The different inhibitory effects on IFN-I induction by two m^6^A-modified RNA oligos compared to their unmodified counterparts might be due to different sequences or conformation of the RNA fragments [32]. We also demonstrated that HIV-1 RNA with decreased m^6^A levels enhanced IFN-I expression, but HIV-1 RNA with increased m^6^A modifications had opposite effects. Our results suggest that HIV-1 genomic RNA and viral transcripts are masked by m^6^A modifications to avoid RIG-I-mediated sensing and IFN-I induction during viral infection. Thus, HIV-1 has likely evolved an immune evasion strategy through m^6^A modification of viral RNA (Fig. S2).

Several RNA modifications, such as N-1-methylpseudouridine, 5-methylcytidine (m^5^C), 5-hydroxymethylcytidine, 5-methoxycytidine, and 2’ fluoro-deoxyribose, have significant impact on RIG-I- and MDA5-mediated RNA sensing [40]. In addition to m^6^A modification, HIV-1 genomic RNA contains eight types of epitranscriptomic modifications that are higher than the average cellular mRNA, with m^5^C and 2’-*O*-methyl modifications being most prevalent [41]. It is possible that HIV-1 RNA exploits multiple epitranscriptomic modifications to avoid innate sensing as mechanisms of immune evasion. This possibility may explain how HIV-1 is able to avoid innate immune responses to establish persistent and latent infection even in patients on combined ART [31].

The IFN-I gene itself is m^6^A-modified and targets its destabilization for the maintenance of homeostatic state in mouse and humans [38]. Rubio *et al*. showed that, following human cytomegalovirus infection, depletion of METTL14 or increase in ALKBH5 proteins leads to decrease level of m^6^A in IFN-β gene and stabilizes and elevates the IFN-I response [37]. In this study, we observed increased m^6^A levels in cellular RNA of PBMCs from HIV-1 viremic patients compared to HIV-1 suppressed patients on ART. However, we did not observe significant changes in the levels of m^6^A writers and erasers in PMBCs from healthy donor, HIV-1 viremic patients, and HIV-1 patients on ART. These results are consistent with our previous data showing increased m^6^A levels in HIV-1 infected primary CD4^+^ T-cells in the absence of altered expression of m^6^A writers or erasers [30]. It is possible that HIV-1 may modulate the activity or localization of writers or eraser, thereby upregulating m^6^A levels in HIV-1 infected cells. It remains to be established whether m^6^A modification of HIV-1 RNA regulate innate immune responses in primary CD4^+^ T-cells or macrophages.

We found that m^6^A-modified HIV-1 reduces the activation of IRF3 and IRF7 through RIG-I-mediated signaling to suppress IFN-I induction. However, it remains unclear how m^6^A modifications of HIV-1 RNA reduces phosphorylation of IRF3 and IRF7 during early stage of HIV-1 infection. Previous studies suggest that HIV-1 proteins can target several cellular RNA and DNA sensors including RIG-I to surpass the IFN-I response [42–45]. Moreover, HIV-1 can also target downstream proteins in the IFN-I pathway including IRF3 and IRF7 to contribute to chronic and persistent infection [46–50]. For example, HIV-1 Vpr protein mediates degradation of IRF3 to avoid the innate antiviral immune response [51].

Durbin *et al*. showed that a RIG-I-activating RNA ligand, the 106-nucleotide polyU/UC sequence derived from the 3’ UTR of hepatitis C virus with m^6^A modification bound RIG-I with low affinity and did not trigger the conversion to the activated RIG-I conformer and thus has an immunosuppressive potential [40]. Our data indicated that m^6^A-defective HIV-1 RNA enhanced RIG-I-mediated RNA sensing and IFN-I induction in cells. Further studies are needed to examine whether the m^6^A-modified HIV-1 RNA binds RIG-I with a low affinity, which might be the possible cause of reduced IFN-I induction during viral infection.

In summary, our study uncovered a previously unidentified strategy of how HIV-1 RNA escapes the host antiviral innate immune system through m^6^A modifications of its RNA genome. HIV-1 RNA m^6^A modifications can act as an immune suppressor of RIG-I-mediated viral RNA sensing. Our findings suggest that pharmacological reduction in m^6^A modification of HIV-1 RNA may enhance IFN-I-mediated innate antiviral immune responses, thereby inhibiting viral replication.

## Materials and Methods

### Cell culture

HEK293T cell line was a kind gift from Vineet KewalRamani (National Cancer Institute, USA) and maintained in complete Dulbecco’s modified Eagle’s medium (DMEM) as described [32]. U937 cell line was obtained from the American Type Culture Collection (ATCC) and maintained in complete RPMI-1640 medium as described [52]. All the cell lines were maintained at 37 °C in 5% CO_2_ and tested negative for mycoplasma contamination using a universal mycoplasma detection kit (ATCC 30-1012K) as described [53].

### Plasmids and HIV-1 RNA oligos

The HIV-1 proviral DNA construct pNL4-3 was used to generate viral stocks as described [19]. For over-expression of the m^6^A erasers, the corresponding control vectors, pCMV6-FTO, pCMV-ALKBH5 were described [9, 54]. For knockout of eraser genes, CRISPR-Cas9 vectors containing sgControl, sgFTO, and sgALKBH5 were used as described [38]. For RIG-I knockout, pCR-BluntII-Topo-sgRIGI-1 and 2 vectors were described [55], which were kindly provided by Dr. Stacy Horner (Duke University, USA), and the plasmid hCas9 (catalog no. 41815, Addgene) was described [56]. For MDA5 and RIG-I knockdown, shControl, shMDA5 and shRIG-I plasmids [29] were kindly provided by Dr. Yamina Bennasser (Université de Montpellier, France). Four RNA oligo sequences are from the 5’ UTR of HIV-1 genomic RNA (NL4-3 strain) with or without a single m^6^A site [32], which were commercially synthesized (Integrated DNA Technologies). The sequences and the location of the m^6^A sites in the conserved <u>GG**A**CU</u> motifs of the HIV-1 genome were described [32] and are listed below: RNA oligo 1 (nt. 235-281, the m^6^A-modified adenosine is nt. 241): 5’-CGCA<u>GG**A**CU</u>CGGCUUGCUG<u>GAG</u>ACGGCAAGAGGCGAGGGGCG-3’. To eliminate RNA dimerization in our previous RNA binding assays [32], the original dimer initiation sequence of HIV-1 (AAGCGCGC) in oligo 1 was replaced with the underlined nucleotides GAG. RNA oligo 2 (nt. 176-217, the m^6^A-modified adenosine is nt. 197): 5’-AGCAGUGGCGCCCGAACAG<u>GG**A**CU</u>UGAAAGCGAAAGUAAAGC-3’.

### Generation of U937 cells with MDA5 knockdown or RIG-I knockout, and HEK293T cells with FTO or ALKBH5 knockout

For MDA5 knockdown U937 cell line construction, HEK293T cells were transfected with shControl or shMDA5, together with pMD2.G and psPAX2 plasmids by polyethyleneimine (PEI) [53]. At 48 h post-transfection, lentiviruses were harvested and purified to infect U937 cells for 48 h and then the U937 cells were selected in RPMI-1640 media with 1 μg/mL puromycin. To generate RIG-I knockout cells, pCR-BluntII-Topo-sgRIGI-I or pCR-BluntII-ToposgRIGI-2, along with hCas9, which has neomycin (G148) resistance, were transfected into U937 cells by TransIT mRNA transfection kit (mirus, USA) for 48 h according to the manufacturer’s protocol. Then, G418 (1 mg/mL) was added to transfected cells for 8 days to select RIG-I knockout U937 cells, which were confirmed by Western blotting. For Control, FTO, and ALKBH5 knockout HEK293T cell generation, HEK293T cells were transfected with corresponding single guide RNAs (sgRNAs), together with pMD2.G and psPAX2 plasmids. At 48 h post-transfection, lentiviruses were collected to infect fresh HEK293T cells for 48 h. Then, the single clones were selected by 1 μg/mL puromycin in 96 well plates. The KO cells were confirmed by DNA sequencing and for specific protein expression by Western blotting.

### Dot immunoblotting of m^6^A modification in RNA

RNA was extracted from purified and concentrated HIV-1 stocks by using TRIzol (Invitrogen) or RNA purification kit (Qiagen). The synthesized RNA oligos were directly used for dot-blot assays as described [30]. Briefly, HIV-1 RNA or RNA oligos (diluted to 100 μL using 1 mM EDTA) were mixed with 60 μL of 20× saline-sodium citrate (SSC) buffer (3 M NaCl, 0.3 M trisodium citrate) and 40 μL of 37% formaldehyde (Invitrogen) and incubated at 65 °C for 30 min. Nitrocellulose membrane (162-0115, Bio-Rad) or nylon membranes (11209299001, Roche) were pre-soaked with 10X SSC for 5 min and assembled in dot-blot apparatus (Bio-Rad) with vacuum-on. Equal amounts of RNA were transferred to nitrocellulose or nylon membranes, then membranes were washed twice with 200 μL of 10× SSC buffer. Nylon membranes were washed once with TBST buffer (20 mM Tris, 0.9% NaCl, and 0.05% Tween 20) for 5 min and stained with methylene blue staining (MB119, Molecular Research Center) for 2-5 sec followed by two or three washes with ddH_2_O. Nitrocellulose membranes were blocked with 5% milk in TBST buffer and used to detect m^6^A levels by probing with m^6^A specific antibodies (Synaptic Systems; 202 003). Images were taken by Amersham Biosciences Imager 600 (GE Healthcare) and analyzed by ImageJ software (National Institutes of Health). Densitometry quantification of relative RNA m^6^A levels was normalized to MB staining as described [30].

### *In vitro* FTO demethylation of HIV-1 RNA m^6^A

Demethylation of HIV-1 RNA m^6^A was performed with recombinant FTO treatment of purified HIV-1 RNA. Briefly, 500 ng HIV-1 RNA were used for FTO *in vitro* treatment in 100 μL reaction buffer containing 50 mM HEPES buffer (pH7.0), 75 μM (NH_4_)_2_Fe (SO_4_)_2_•6H_2_O, 2 mM L-ascorbic acid, 300 μM L-ascorbic acid, 200U RNAsin, 5 μg/mL BSA, and 0.2 nmol FTO protein. The reaction was performed at 37 °C for 1 hr and then stopped by adding 5 mM EDTA. Finally, RNA samples were denatured at 70 °C for 2 min and quickly put into ice for m^6^A detection.

### HIV-1 production, p24 quantification, U937 cells transfection and HIV-1 infection assays

HIV-1 stocks were generated by transfection of HEK293T cells with the proviral DNA pNL4-3 using PEI as described [53]. Cell culture medium was exchanged at 6-8 h post-transfection with supernatants and was harvested at 48 h. The cell culture media containing viruses were filtered (0.45 μm) and purified by 25% sucrose using an SW28 rotor (Beckman Coulter) at 141,000*g* for 90 min. The pellet was resuspended with PBS and digested with DNase I (Turbo, Invitrogen) for 30 min at 37 °C. To extract HIV-1 genome RNA, concentrated HIV-1 virions were lysed by Trizol (Invitrogen) and RNA was purified by phenolic-chloroform sedimentation and isopropanol precipitation. For transfection, cells were treated with 100 ng/mL phorbol 12-myristate 13-acetate (PMA) for 24 h and changed with fresh RPMI-1640 media for another 24 h. PMA-differentiated U937 cells were then transfected with TransIT mRNA transfection kits (Mirus) according to the manufacturer protocol. At 16 h post-transfection, cells were harvest for RT-qPCR analysis. For infection assays, HIV-1 p24 levels were quantified by an enzyme-linked immunosorbent assay (ELISA) using anti-p24-coated plates (The AIDS and Cancer Virus Program, NCI-Frederick, MD) as described [30]. PMA-differentiated U937 cells were infected by equal amounts of HIV-1 (250 pg of p24) for 16 h and then cells were collected for Western blotting or RT-qPCR analysis.

### Antibodies and immunoblotting

The antibodies used in this study were: anti-GAPDH (AHP1628, Bio-Rad), anti-FLAG (F1804, Sigma-Aldrich), anti-METTL3 (15073-1-AP, Proteintech Group), anti-METTL14 (HPA038002, Sigma-Aldrich), anti-FTO (ab124892, Abcam), anti-ALKBH5 (HPA007196, Sigma-Aldrich), anti-MDA5 (D74E4, Cell signaling), anti-RIG-I (D14G6, Cell signaling), anti-HIV-1 Gag (clone #24-2, the NIH AIDS Reagent Program), anti-IRF3 (124399, Abcam), anti-phospho-IRF3 (49475, Cell Signaling), anti-IRF7 (SC-9083, Santa Cruz), anti-phospho-IRF7 (5184, Cell Signaling) and anti-m^6^A polyclonal rabbit Ab (202003, Synaptic Systems). Cells were harvested and lysed in cell lysis buffer (Cell Signaling) supplemented with protease inhibitor cocktails (Sigma-Aldrich). Immunoblotting was performed as described [30]. Detection of GAPDH expression was used as a loading control.

### Quantitative RT-PCR

Real-time quantitative RT-PCR (qRT-PCR) was performed as described [53] to assess the relative levels of *IFN-α* and *IFN-β* mRNA expression in cells induced by HIV-1 RNA transfection or HIV-1 infection. Following primers (IDT) were used:

*IFN-α*, F 5’-GTACTGCAGAATCTCTCCTTTCTCCT-3’

*IFN-α*, R 5’-GTGTCTAGATCTGACAACCTCCCAGG-3’

*IFN-β*, F 5’-AACTTTGACATCCCTGAGGAGATTAAGC-3’

*IFN-β*, R 5’-GACTATGGTCCAGGCACAGTGACTGTAC-3’

*GAPDH*, F 5’-GGAAGGTGAAGGTCGGAGTCAACGG-3’

*GAPDH*, R 5’-CTGTTGTCATACTTCTCATGGTTCAC-3’

### Ethics statement

The study using human PBMCs from healthy control subjects and HIV-positive individuals has been approved by the Institutional Review Board of the University of Iowa. The study was conducted according to the Declaration of Helsinki guidelines.

### PBMCs from healthy donors and HIV-1 patients

Healthy control subjects and HIV-positive individuals attending the University of Iowa HIV Clinic who were receiving ART and had HIV-1 viral load levels below the limit of detection (< 20 copies/mL) for over 6 months were invited to participate in these studies, and all provided written informed consent. HIV-1 viral load was determined using the COBAS^®^ AmpliPrep/COBAS^®^ TaqMan HIV-1 test (Roche). PBMCs were purified using BD Vacutainer^®^ CPT™ Mononuclear cell preparation tubes (BD Biosciences) as recommended by the manufacturer. Cells were stored in 92% fetal calf serum, DMSO in liquid nitrogen until use. PBMCs were obtained with 9 healthy donors, 6 HIV-1 viremic patients, and 16 HIV-1 patients treated with ART (Supplemental Table 1). Both viral RNA and protein were isolated from these PBMCs at the same day using Ambion Paris RNA and protein extraction kit (ThermoFisher Scientific) and stored in −80 °C until use. The RNA was quantitated using the NanoDrop spectrophotometer (ThermoFisher Scientific) and was used for m^6^A dot-blot detection (200 ng) and *IFN-α* and *IFN-β* mRNA analyses as described [53]. The protein was quantitated using Pierce BCA reagent (ThermoFisher Scientific) and subjected to Western blot analysis of the m^6^A writers and erasers.

### Statistical analyses

Data were analyzed using either Mann-Whitney’s t-test or the one-way analysis of variance (ANOVA) with Prism software and statistical significance was defined as *P* < 0.05. All experiments were repeated at least three times.

## Acknowledgments

This work was supported in part by National Institutes of Health grants R01AI150343 and R01AI141495 (to L. W.). We thank Alexis Hawkins for technical assistance in detection cellular m^6^A levels of PBMCs, and the Wu lab members for helpful discussions and suggestions. We also thank Michael Cahill for outstanding technical support. The authors appreciate generous reagents from Drs. Yamina Bennasser, Stacy Horner, Vineet KewalRamani, Noam Stern-Ginossar, and the NIH AIDS Reagent Program.

## Author contributions

S.C., S.K., and N.T. performed experiments and contributed to manuscript preparation. J.W. and J.T.S. provided PBMCs from HIV-1 patients and uninfected individuals, and helped data analyses. L.H. and C.H. provided recombinant FTO and the *in vitro* m^6^A demethylation protocol. All authors analyzed data and contributed to experiment design. S.C. and S.K. drafted the manuscript. L.W. conceived the study, supervised the work, and revised the manuscript. All authors contributed to manuscript editing and revision.

## Competing interests

The authors declare no competing interests.

## Supplemental Fig. S1-S2 legends

**Fig. S1.**
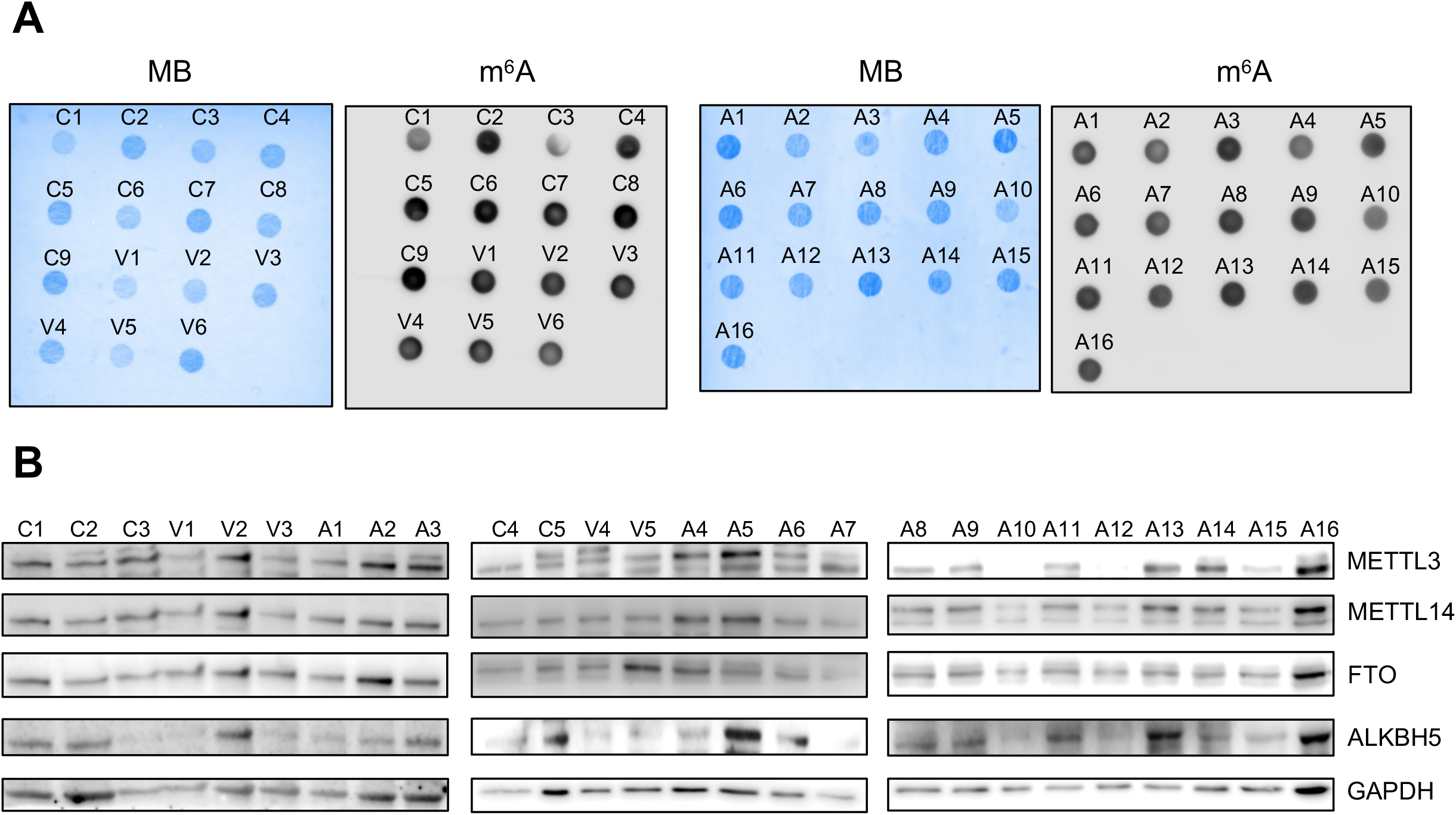
Detection of RNA m^6^A modification and writer and eraser proteins in PBMCs from healthy donors and HIV-1 patients. **(A)** Total cellular RNA was isolated from the PBMCs of three groups; uninfected healthy control individuals (C1-C9), HIV-1 infected individuals without ART (V1-V6) and with ART (A1-A16) were subjected to m^6^A dot-blot analysis (200 ng RNA/sample). Methylene blue (MB) staining serves as a loading control. Here every blot represents one patient with the code as referred in Supplemental Table 1. **(B)** Cell lysates of PBMCs of each group uninfected healthy individuals (C1-C5), HIV-1 infected individuals without ART (V1-V5) and with ART (A1-A16) were subjected to Western blot analysis. Five samples (C6-C9 and V6) were not included in the Western blot analysis due to the lack of sufficient cell lysates. Equal amount of proteins (10 μg) of whole cell lysate was immunoblotted for m^6^A writers (METTL3 and METTL14) and erasers (ALKBH5 and FTO) using specific antibodies. GAPDH serves as a loading control. For the densitometry quantitation of METTL3 levels, only one band at an approximate molecular weight of 70 kDa was used.

**Fig. S2.**
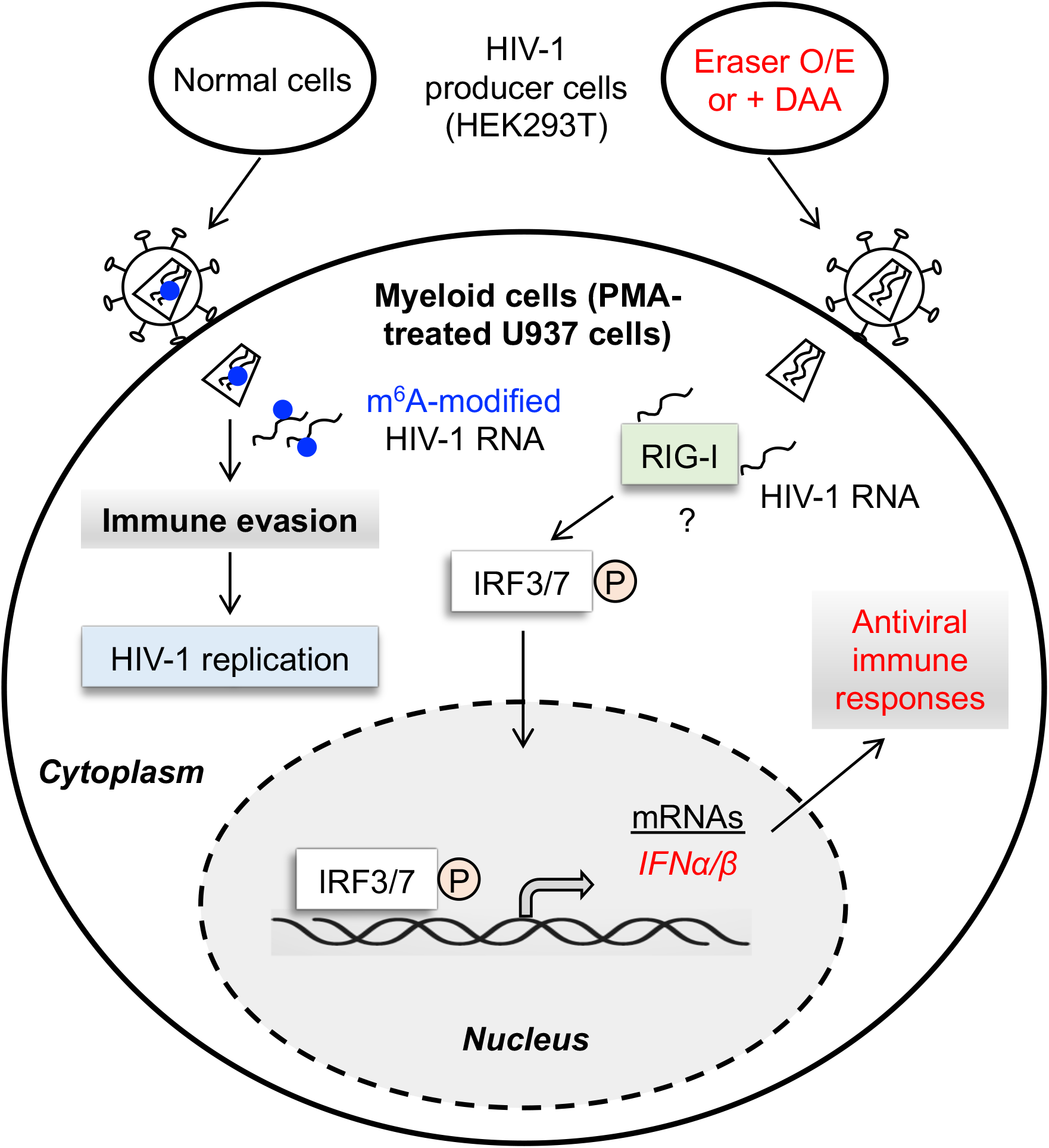
HIV-1 RNA escapes from innate immune surveillance. In HIV-1 producer cells, writers add and erasers remove internal m^6^A modifications (blue dots) of viral RNA, respectively. HIV-1 with m^6^A-modificed RNA avoids innate sensing in infected myeloid cells, thereby escaping immune surveillance. Overexpression (O/E) of erasers or inhibiting m^6^A addition with DAA in HIV-1 producer cells generates viruses with m^6^A-defective viral RNA. When HIV-1 with m^6^A-defective RNA infects macrophage-like cells, the cytoplasmic RNA sensor RIG-I recognizes unmodified HIV-1 RNA and triggers phosphorylation (indicated by the letter P) of the transcription factors IRF3 and IRF7. Phosphorylation of IRF3/7 leads to IFN-α/β expression and generates antiviral innate immune responses in HIV-1-infected macrophage-like cells. However, it remains to be established whether m^6^A-defective HIV-1 RNA enhances binding to RIG-I, thereby inducing IRF3/7 activation and IFN-I expression in cells.

**Supplemental Table 1.** Details of PBMC samples from healthy control donors (C1-C9), HIV-1 viremic patients (V1-V6), and HIV-1 patients on ART (A1-A16)

| **Sample code** | **Donor gender** | **Infection status** | **HIV-1 viral load (copies/mL)** | **Therapy regimen** |
| --- | --- | --- | --- | --- |
| C1 | Male (M) | Uninfected | 0 | No therapy |
| C2 | M | Uninfected | 0 | No therapy |
| C3 | M | Uninfected | 0 | No therapy |
| C4 | M | Uninfected | 0 | No therapy |
| C5 | M | Uninfected | 0 | No therapy |
| C6 | M | Uninfected | 0 | No therapy |
| C7 | M | Uninfected | 0 | No therapy |
| C8 | M | Uninfected | 0 | No therapy |
| C9 | M | Uninfected | 0 | No therapy |
| V1 | M | HIV+Viremic | 120,000 | Pre-therapy |
| V2 | M | HIV+Viremic | 34,000 | Pre-therapy |
| V3 | M | HIV+Viremic | 1,318,000 | Pre-therapy |
| V4 | M | HIV+Viremic | 35,000 | Pre-therapy |
| V5 | M | HIV+Viremic | 16,000 | Pre-therapy |
| V6 | M | HIV+Viremic | 22,909 | Pre-therapy |
| A1 | M | HIV+Suppressed | ND (non-detectable) | Abacavir, Lamivudine, Efavirenz |
| A2 | M | HIV+Suppressed | ND | Tenofovir, emtricitabine, dolutegravir |
| A3 | M | HIV+Suppressed | ND | Abacavir, Lamivudine, dolutegravir |
| A4 | M | HIV+Suppressed | ND | Tenofovir, emtricitabine, dolutegravir |
| A5 | M | HIV+Suppressed | ND | Abacavir, Lamivudine, dolutegravir |
| A6 | M | HIV+Suppressed | ND | Tenofovir,<br>emtricitabine, Efavirenz |
| A7 | M | HIV+Suppressed | ND | Tenofovir,<br>emtricitabine,<br>dolutegravir |
| A8 | M | HIV+Suppressed | ND | Tenofovir,<br>emtricitabine,<br>dolutegravir |
| A9 | M | HIV+Suppressed | ND | Tenofovir,<br>emtricitabine, Efavirenz |
| A10 | M | HIV+Suppressed | ND | Abacavir, Lamivudine,<br>dolutegravir |
| A11 | M | HIV+Suppressed | ND | Abacavir, Lamivudine,<br>dolutegravir |
| A12 | M | HIV+Suppressed | ND | Abacavir, Lamivudine,<br>Efavirenz |
| A13 | M | HIV+Suppressed | ND | Tenofovir,<br>emtricitabine,<br>elvitegravir, cobicistat |
| A14 | M | HIV+Suppressed | ND | Abacavir, Lamivudine,<br>dolutegravir |
| A15 | M | HIV+Suppressed | ND | Abacavir, Lamivudine,<br>dolutegravir |
| A16 | M | HIV+Suppressed | ND | Abacavir, Lamivudine,<br>dolutegravir |

## Notes

### Competing Interest Statement

The authors have declared no competing interest.

